# The small molecule KHS101 induces bioenergetic dysfunction in glioblastoma cells through inhibition of mitochondrial HSPD1

**DOI:** 10.1101/205203

**Authors:** Euan S. Polson, Verena B. Kuchler, Christopher Abbosh, Edith M. Ross, Ryan K. Mathew, Hester A. Beard, Eulashini Chuntharpursat-Bon, Jennifer Williams, Bárbara Da Silva, Hao Shao, Anjana Patel, Adam J. Davies, Alastair Droop, Hollie B.S. Griffiths, Paul Chumas, Susan C. Short, Mihaela Lorger, Jason Gestwicki, Lee D. Roberts, Robin S. Bon, Simon J. Allison, Shoutian Zhu, Florian Markowetz, Heiko Wurdak

## Abstract

Pharmacological inhibition of uncontrolled cell growth with small molecule inhibitors is a potential strategy against glioblastoma multiforme (GBM), the most malignant primary brain cancer. Phenotypic profiling of the neurogenic small molecule KHS101 revealed the chemical induction of lethal cellular degradation in molecularly-diverse GBM cells, independent of their tumor subtype, whereas non-cancerous brain cells remained viable. Mechanism-of-action (MOA) studies showed that KHS101 specifically bound and inhibited the mitochondrial chaperone HSPD1. In GBM but not non-cancerous brain cells, KHS101 elicited the aggregation of an enzymatic network that regulates energy metabolism. Compromised glycolysis and oxidative phosphorylation (OXPHOS) resulted in the metabolic energy depletion in KHS101-treated GBM cells. Consistently, KHS101 induced key mitochondrial unfolded protein response factor DDIT3 *in vitro* and *in vivo*, and significantly reduced intracranial GBM xenograft tumor growth upon systemic administration, without discernible side effects. These findings suggest targeting of HSPD1-dependent oncometabolic pathways as an anti-GBM therapy.

## Introduction

GBM is the most common malignant primary brain tumor in adults and among the most devastating cancers (1). Its overall median time to recurrence after surgery and standard chemoradiotherapy is ^~^7 months and the 5-year survival rate remains low (<5%) (2). As examples of the potential therapeutic promise shown by small molecules, pre-clinical data support anti-GBM effects through perturbation of cell death programs (3), transcriptional and epigenetic pathways (4, 5), lethal autophagy (6), and GBM stem cell self-renewal (7). However, GBM biology remains poorly understood and there is an unmet need for the identification of molecular GBM vulnerabilities and the development of pharmacological strategies to exploit them (2). In addition, to yield potential novel drug candidates, phenotypic drug discovery and profiling approaches can address the complexity of diseases, in particular, when the molecular target(s) and MOA that underlie a small molecule effect are identified (8, 9).

Here, we hypothesized that an experimental small molecule could identify molecular vulnerabilities in patient-derived GBM cells that possess stem cell-like characteristics (10). Our investigation focused on the 4-aminothiazole derivative KHS101 as it penetrates the blood brain barrier (BBB) and promotes neural progenitor cell differentiation at the expense of proliferation in the adult rat (11). As inter-and intratumor heterogeneity is a major impediment to broadly efficacious GBM therapy, we sought to address whether KHS101 can affect a spectrum of clinically-relevant GBM subtype(s). To achieve this, we established a panel of different patient-derived primary and recurrent GBM cell models that were characterized through cytogenetic and single cell gene expression analysis. We observed that KHS101 leads to a rapid and selective cytotoxic response in this heterogeneous spectrum of patient-derived GBM cell models. Accordingly, we sought to identify the MOA behind the KHS101 anti-GBM activity utilizing affinity-based target identification, orthogonal chemical validation, and detailed analysis of energy metabolism and mitochondrial proteostasis. Furthermore, we investigated the compound's anti-GBM activity in established xenograft tumors upon systemic administration.

## Results

### KHS101 selectively induces GBM cell cytotoxicity independent of molecular subtypes

GBM is characterized by intra-and intertumor heterogeneity that may hinder therapeutic success (12–14). To represent this molecular heterogeneity, we established six patient-derived tumor cell models from primary GBM (GBM1, 4, 13), recurrent GBM (GBM20), and rare GBM subtypes such as gliosarcoma (GBM11) and recurrent giant cell GBM (GBM14, Fig. 1A). Molecular subtype profiling was carried out through single cell qRT-PCR in ^~^45 randomly selected cells from each tumor model. Based on the differential expression of 88 ‘stemness’, cell cycle, and GBM subtype classifier genes, computational analysis revealed that our GBM models contain heterogeneous cell populations with single (mesenchymal or proneural) or double (classical/proneural, mesenchymal/proneural) subtype signatures (Fig. 1B and C, Supplementary Report 1). Independent of clinical and molecular subtype features (including classical, proneural, and mesenchymal subtype compartments, primary versus recurrent tumor origin, and MGMT hypo-and hypermethylation), KHS101 attenuated cell growth and survival in several different GBM cell models but had no discernible effect on non-cancerous adult brain progenitor (NP) cells (whose derivation is described in (10); Fig. 1D). Dose response curves further indicated selective cytotoxicity of KHS101 against all GBM cell models as well as GBM cell lines U251 and U87, while two different NP lines (NP1 and NP2) were refractory (Fig. 1E, Supplementary Fig. 1). Consistent with a KHS101-induced pro-apoptotic cell fate, a marked increase in caspase 3/7 activation, increased expression of the harakiri activator of apoptosis (HRK), and accumulation of Annexin V-positive apoptotic cells was observed 48 hours after KHS101 addition (Fig. 1F, G, and H). Electron microscopy (EM) imaging performed 12 hours after KHS101 addition revealed that GBM apoptosis was preceded by the pronounced development of intracellular vacuoles and degradation of electron-dense cytoplasmic content in GBM compared with NP cells (Fig. 2A and B, Supplementary Fig. 2). Concomitantly, we observed an increase in the LC3B-positive autophagosomal compartment in all GBM but not NP cell models (Fig. 2C, D, and E). Neither bone morphogenetic protein 4 (BMP4)-induced differentiation of GBM cells (15) nor reduced oxygen tension (5% O_2_) (16) modulated KHS101 cytotoxicity (Fig. 2F and G). Taken together, these findings indicate a selective induction of GBM cell cytotoxicity by a single agent in a spectrum of disease-relevant cellular contexts.

**Fig. 1.**
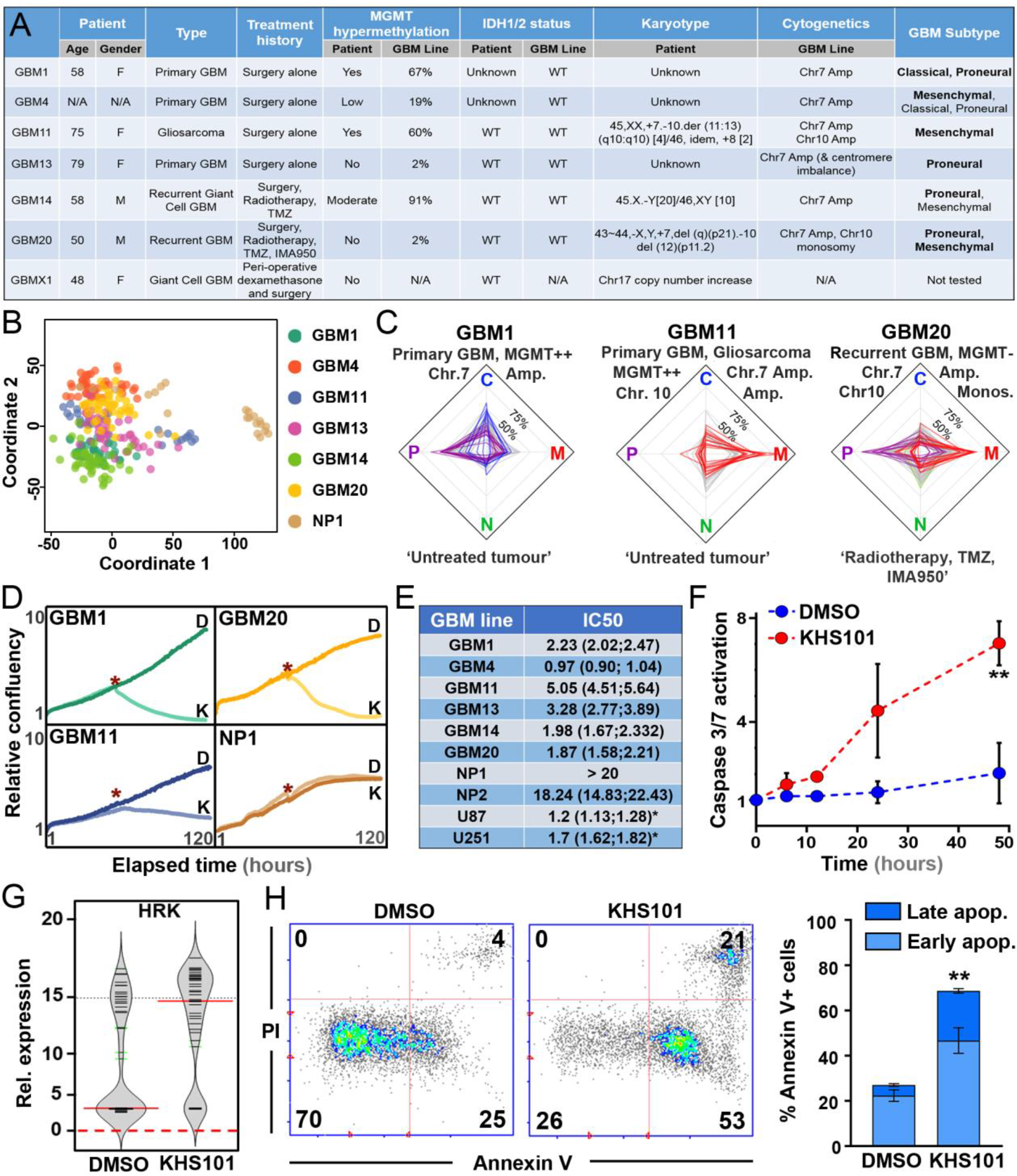
KHS101 exhibits cytotoxicity in molecularly-diverse GBM models. (A) Overview of patient-derived GBM cell model characterization. (B) Principle components analysis of single cells derived from six patient-derived GBM lines and the NP1 model. (C) Radar plots depicting the GBM subtype compartments (classical (C), proneural (P), mesenchymal (M) and neural (N)) alongside clinical and cytogenetic features of GBM1 (left), GBM11 (middle) and GBM20 (right) models. (D) Time-lapse microscopy-based quantification of cell growth rates (GR, % per hour) of GBM1 (0.019%), GBM11 (0.012%), GBM0 (0.015%) and NP1 (0.018%). GRs decline in GBM1 (-0.018%), GBM11 (-0.01%), GBM20 (-0.018%) but not NP1 cells upon addition of KHS101 (7.5 μM, K, red asterisks indicate start of treatment) compared to the DMSO control (0.1%, D). (E) Five day IC50 values of KHS101-dependent cell viability (and 95% confidence intervals) are shown for all tested GBM, NP and glioma cell lines. Asterisks indicate 48 hour IC50 values. (F) Increase in caspase 3/7 activation in GBM1 cells treated with KHS101 (7.5 μM) compared with DMSO (0.1%). Data were normalized to t_0_. (G) Single cell qRT-PCR-based bean plot indicating upregulation of the activator of apoptosis gene harakiri (HRK) in GBM1 cells treated with DMSO (0.1%) or KHS101 (7.5 μM) for 24 hours. Solid red bars indicate mean expression values, black lines represent expression values of the individual cells, and dashed red lines separate expressors from non-expressors. (H) Flow cytometry data analysis indicates an increased number of early apoptotic (Annexin V-positive/PI-negative) and late apoptotic (Annexin V-positive/PI-positive) GBM1 cells 48 hours after KHS101 (7.5 μM, right) compared with DMSO treatment (0.1%, left). Data are mean ± SD of 3 biological replicates. **, *P*<0.01, student’s t-test (two tailed).

**Fig.2.**
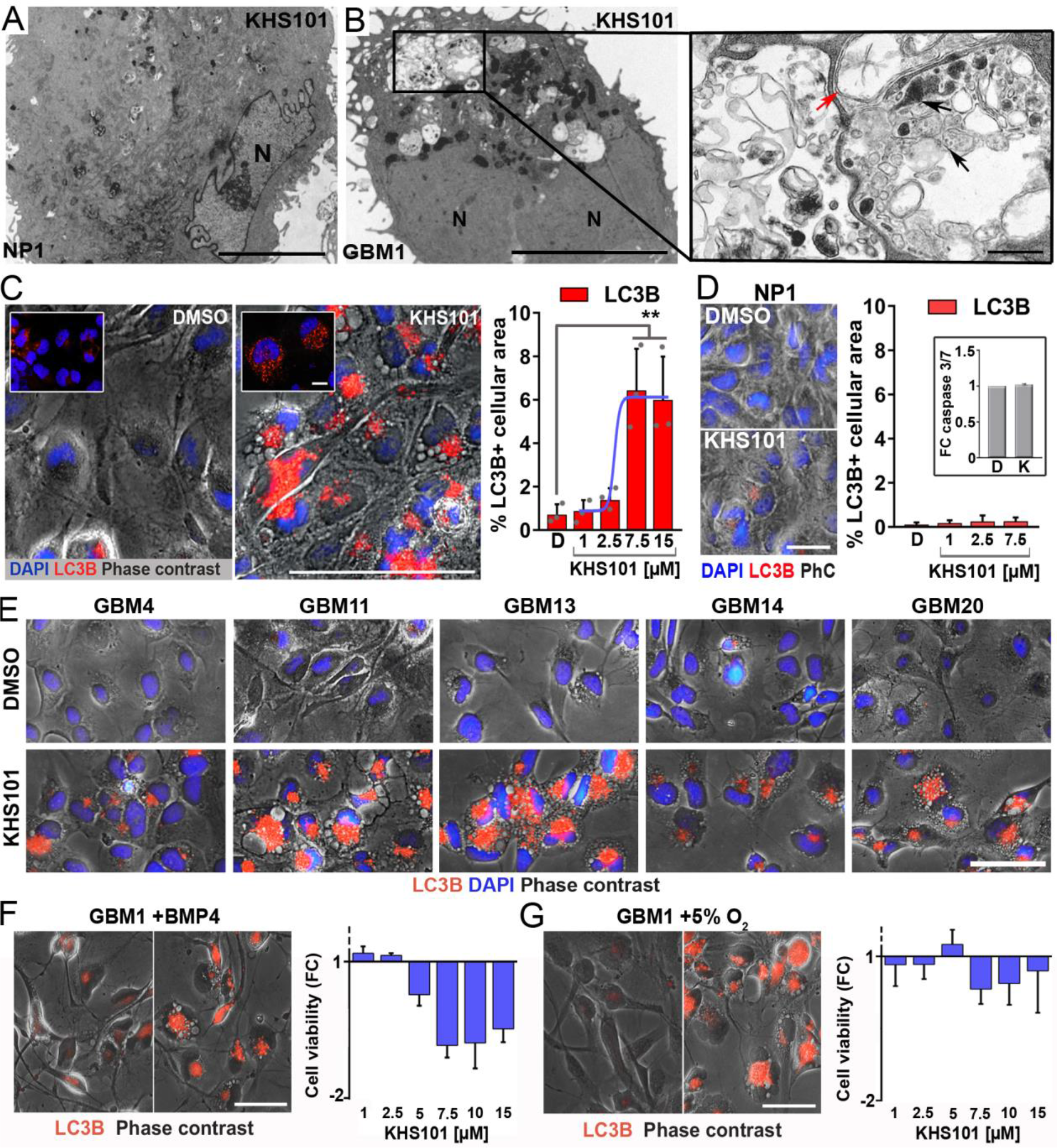
KHS101 exhibits cellular degradation and autophagy in GBM cells. (A and B) EM images of NP (A) and mitotic GBM cell (B) 12 hours after KHS101 treatment (7.5 μM). Higher magnification image shows the vacuolated area within the KHS101-treated GBM cell highlighting double-membraned autophagosomes (red arrow) alongside larger vacuoles containing cellular material at varying degrees of degradation (black arrows). Scale bars, 5 μm, 500 nm, ‘N’ indicates nuclei. (C) Immunofluorescence images and quantification (EC50 = 2.9 ± 0.84 μM) of LC3B staining in GBM1 cells treated with DMSO (0.1%) or KHS101 (7.5 μM) for 12 hours. Inlays highlight punctate LC3B localization. Scale bars, 100 μm, 10 μm. (D) Immunofluorescence images and quantification of LC3B staining in NP cells treated with DMSO (D; 0.1%) or KHS101 for 12 hours. PhC: phase contrast. Scale bar, 25 μm. Inlay graph shows caspase 3/7 activation fold changes (FC) in DMSO (D; 0.1%) and KHS101-treated (K; 7.5 μM) NP1 cells. (E) LC3B immunofluorescence images of GBM4, GBM11, GBM13, GBM14 and GBM20 cells treated for 12 hours with DMSO (0.1%) or KHS101 (7.5 μM). Images reveal increase in LC3B punctate staining in all GBM cell types after KHS101 as compared with DMSO treatment. Scale bar, 50 μm. (F) Immunofluorescence images of GBM1 cells treated with BMP4 (100 ng/mL). Four days after differentiation-promoting treatment, addition of KHS101 (7.5 μM) induces the accumulation of cytoplasmic LC3B compared to the DMSO control (0.1%) at the 12 hour time point. Scale bar, 50 μm. Differentiated versus undifferentiated GBM1 cell viability in response to KHS101 treatment (after 48 hours, at the indicated concentrations) is shown as fold change (FC). KHS101-induced GBM cell cytotoxicity is only marginally reduced by BMP4 differentiation. (G) LC3B immunofluorescence images of GBM1 cells treated for 12 hours with DMSO (0.1%) or KHS101 (7.5 μM) in 5% oxygen. KHS101 treated cells show accumulation of cytoplasmic LC3B. Scale bar, 50 μm. KHS101-induced GBM cell cytotoxicity is equivalent in 5% versus 21% oxygen culture conditions. Data are mean ± SD of 3 biological replicates (shown as individual dots in *C*), **, P<0.01, student’s t-test (two tailed).

### KHS101 binds mitochondrial HSPD1 in GBM cells

TACC3 is a relevant target of KHS101 in rodent neural progenitor cells (11), and KHS101 has been shown to cause cellular destabilization of TACC3 (17). Consistently, Western blot analysis showed that KHS101 reduced TACC3 levels from ^~^18 hours onwards in the GBM1 model (comparable to *TACC3* knockdown, Supplementary Fig. 3A and B). However, a rapidly-increasing autophagic response was observed in the KHS101-treated GBM cells before the levels of TACC3 were reduced by the compound (Fig. 3A). Moreover, the cytotoxic KHS101 phenotype was not recapitulated by TACC3 knockdown, hence excluding TACC3 downregulation as the relevant MOA (Supplementary Fig. 3C). Microarray transcriptome profiling (ArrayExpress, accession E-MTAB-5713) and gene set enrichment analysis (GSEA) of GBM1 cells in the presence of KHS101 suggested that, in addition to the cell cycle, oxidative phosphorylation (OXPHOS) and the tricarboxylic acid (TCA) cycle were significantly altered by KHS101 (Fig. 3B). Consistently, extracellular flux analysis (protocols described in(18)) revealed an acute decline in basal oxygen consumption rates (OCR; ^~^40%) and mitochondrial oxidative capacity (by ≥70%) in GBM1 cells (Fig. 3C).

**Fig. 3.**
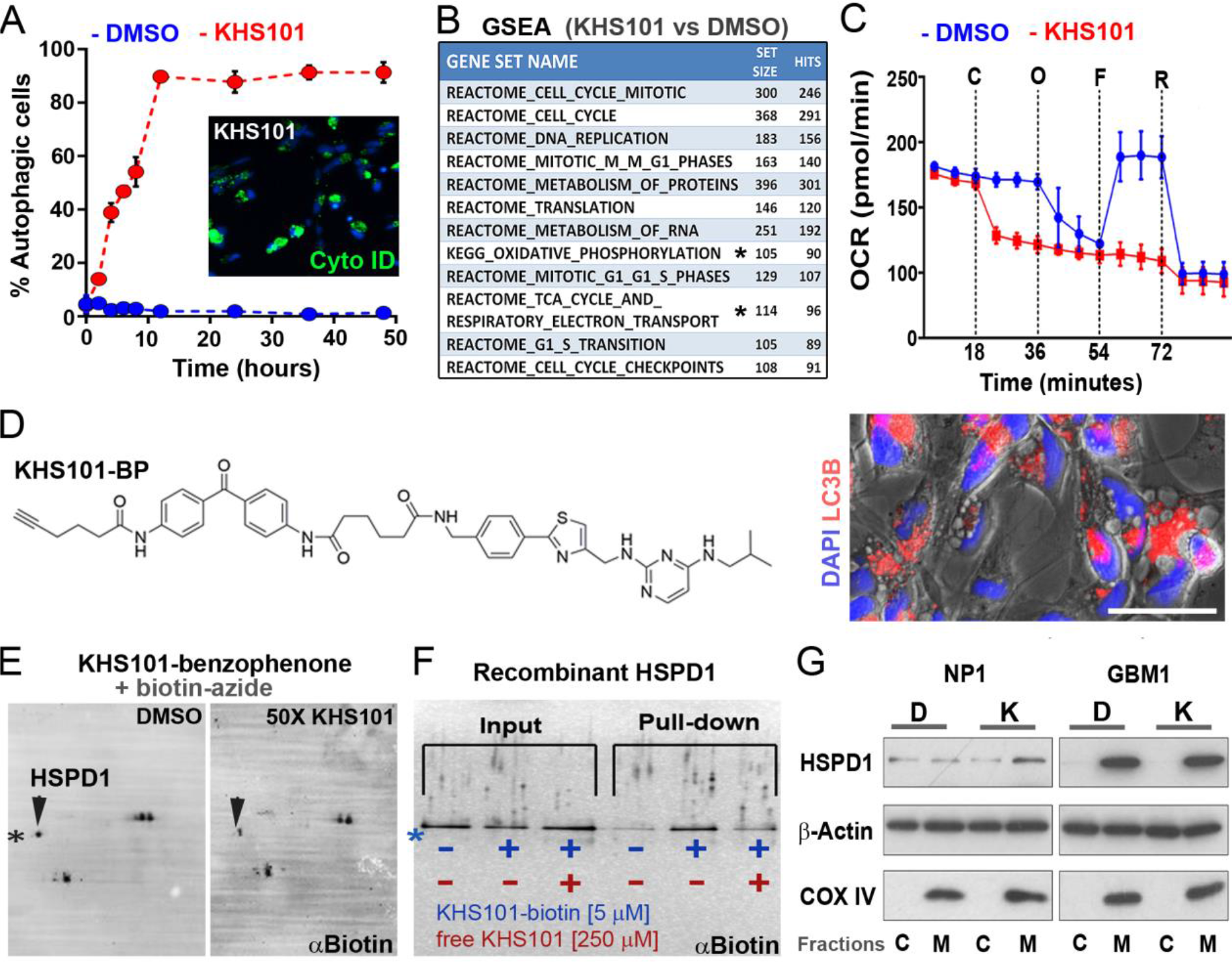
KHS101 obstructs OXPHOS and binds HSPD1. (A) The percentage of autophagic (CytoID^®^-positive) GBM1 cells increases rapidly upon KHS101 (7.5 μM; inlay) vs. DMSO (0.1%) addition. (B) Gene set enrichment analysis indicating major alterations in cell cycle/mitosis and metabolic pathways (including OXPHOS and the TCA cycle marked by asterisks) in GBM1 cells after 24 hours of KHS101 treatment (7.5 μM). (C) GBM1 stress profile of mitochondrial respiration upon addition of KHS101 (7.5 μM) compared with DMSO (0.1%). C: compound addition (DMSO or KHS101), O: oligomycin (1 μM), F: FCCP (0.5 μM), R: roteneone/antimycin A (0.5 μM). (D) Chemical structure of KHS101-Benzophenone (KHS101-BP) and representative LC3B immunofluorescence image of GBM1 cells treated with KHS101-BP (7.5 μM) for 12 hours. Scale bar: 30 μm. (E) Two-dimensional SDS/PAGE and Western blotting of GBM1 cell lysates (20-40% ammonium sulfate-precipitated fraction) detecting KHS101-BP-labeled proteins after photocrosslinking (30 minutes) and biotin-tag labeling (click chemistry with biotin-azide). Arrow points to distinct ^~^60-kDa protein that was reproducibly competed away by free KHS101 and corresponded to HSPD1 as indicated by proteomics analysis. (F) Specific binding of recombinant human HSPD1 with biotinylated KHS101 (in buffer) was detected by silver staining of SDS/PAGE gels in presence/absence of free compound, precipitated with streptavidin-conjugated agarose beads. (G) Immunoblot detecting HSPD1 in GBM1 compared with NP1 cells. HSPD1 levels remain comparable 6 hours after DMSO (0.1%, D) or KHS101 (7.5 μM, K) treatment. Antibodies against β-Actin and COX IV were used loading and mitochondrial/cytoplasmic separation controls. Data are mean ± SD of 3 biological replicates.

Dysregulated mitochondrial processes are important mediators of tumorigenesis (19). To elucidate the cellular target(s) underlying the reduced mitochondrial capacity in KHS101-treated GBM cells, we investigated the physical interaction of KHS101 with potential cellular protein(s) using an established affinity-based target identification protocol (11). The photoaffinity probe KHS101-BP (a KHS101 derivative containing a benzophenone moiety and an alkyne substituent) and KHS101 showed similar bioactivity in GBM cells (Fig. 3D, Supplementary Fig. 4A and B). A distinct KHS101-BP-protein complex of ^~^60 kDa (pI^~^5.7), that was competed away with a 50-fold excess of free KHS101, was identified (Fig. 3E). Proteomics analysis revealed that KHS101-BP-bound protein corresponded to the mitochondrial 60 kDa heat shock protein 1 (HSPD1). KHS101-HSPD1 binding was confirmed in pull-down and thermal shift experiments *in vitro* using human recombinant HSPD1 protein (Fig. 3F and Supplementary Fig. 4C). Cellular fractionation followed by Western blotting showed that HSPD1 was overexpressed in GBM cells and predominantly localized to the mitochondria (Fig. 3G). Consistent with a reported role for HSPD1 in proliferation and complex I integrity in the glioma cell line U87(20), we found that stable HSPD1 knockdown led to a decline in both proliferation and mitochondrial oxidative capacity in GBM1 cells, suggesting an increased dependency on HSPD1 expression and function (Supplementary Fig. 5). However, KHS101 altered neither *HSPD1* expression nor HSPD1 protein levels (Fig. 3G), hence excluding knockdown kinetics as a biologically relevant mechanism underlying the KHS101-GBM cytotoxicity.

### KHS101 selectively aggregates HSPD1 and metabolic enzymes in GBM cells promoting their metabolic exhaustion

To investigate whether the KHS101-HSPD1 interaction inhibits HSPD1 enzymatic function, we assessed HSPD1 chaperone activity *in vitro* upon KHS101 addition. KHS101 elicited a concentration-dependent inhibition of substrate re-folding (Fig. 4A). Furthermore, the GBM cell degradation phenotype and OXPHOS reduction elicited by KHS101 was recapitulated by the mitochondrial HSPD1-binding natural product myrtucommulone (MC (21); Fig. 4B and Supplementary Fig. 6). As these data further confirmed the HSPD1-targeting properties of KHS101, we quantified protein aggregation by fractionation of detergent-insoluble mitochondrial proteins in GBM compared with NP control cells. Silver staining indicated that aggregated proteins were selectively enriched in GBM1 compared with NP1 cells 1 hour after KHS101 addition (Fig. 4C). Proteomics analysis identified HSPD1 as well as enzymes with functions in glycolysis (e.g., ALDOA), TCA cycle (e.g. DLST), oxidative phosphorylation (OXPHOS; e.g. ATP5A1), and mitochondrial integrity (e.g., LONP1 (22)) in the aggregated fractions specifically in the KHS101-treated GBM cells (Supplementary Table 1). These proteins readily integrated into a predicted functional protein-protein interaction network (Fig. 4D). Consistent with the dysfunction of GBM cell metabolism-regulating enzymes, qRT-PCR analysis of 25 key genes altered in the GBM1 transcriptome analysis (ArrayExpress, accession E-MTAB-5713) displayed a KHS101-induced signature of glycolytic/ oxidative/autophagic stress accompanied by loss of known GBM ‘stemness’ markers (e.g., NOS2 (23), ID1 (24), and OLIG2 (25)) in KHS101-treated GBM versus NP cells (Fig. 4E).

**Fig. 4.**
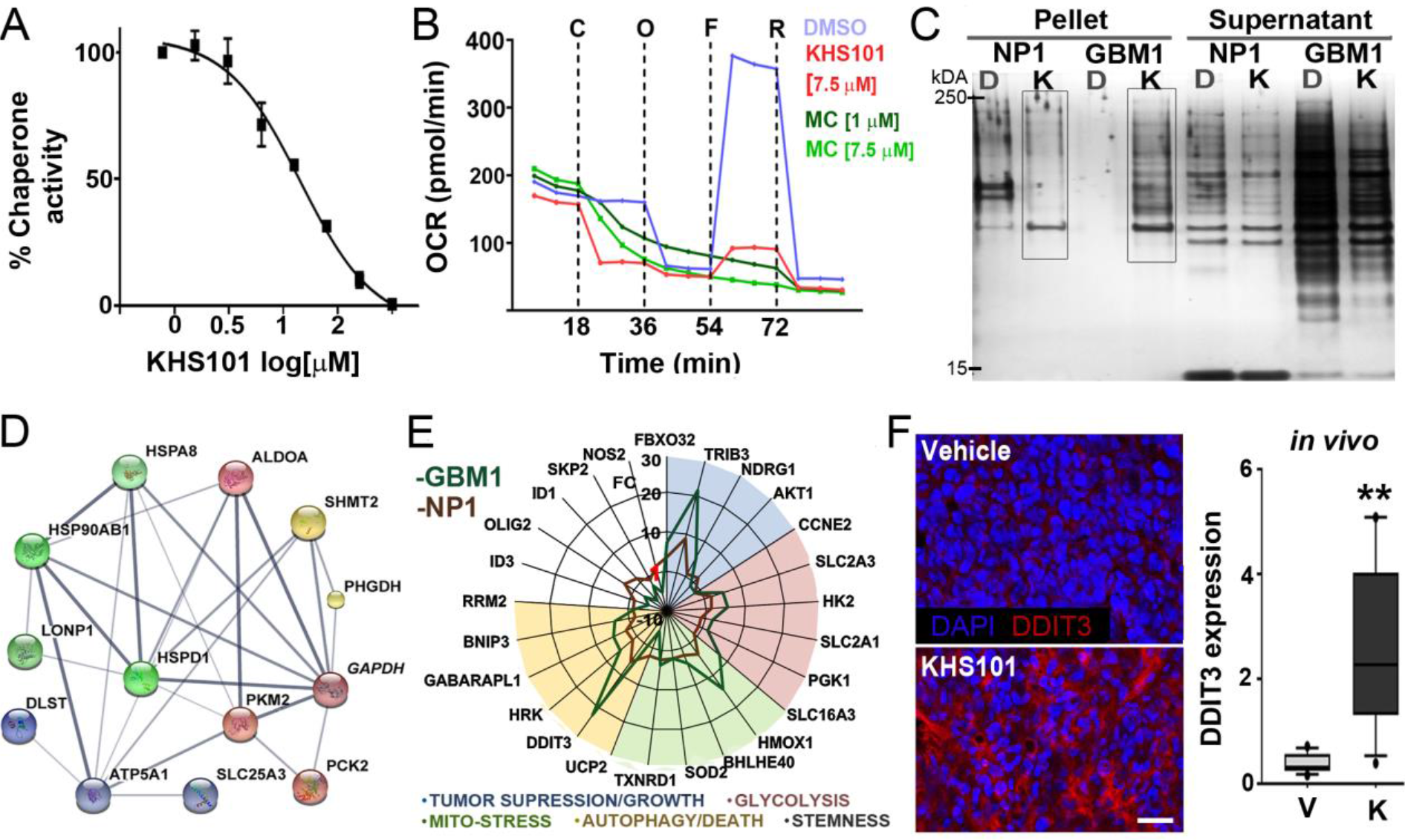
KHS101 promotes selective metabolic enzyme aggregation in GBM cells. (A) Concentration-dependent inhibition of HSPD1 complex substrate refolding activity by KHS101 (IC50 = 14.4 μM). (B) MC reduces mitochondrial capacity in GBM1 cells compared to the control (DMSO) treatment. C: compound addition (DMSO, KHS101 or MC at indicated concentrations), O: oligomycin (1 μM), F: FCCP (0.5 μM), R: roteneone/antimycin A (0.5 μM) (C) Silver staining of NP40 (0.5%) solubilized aggregated (pellet) and soluble (supernatant) mitochondrial fractions from NP1 and GBM1 cells treated with DMSO (0.1%, D) or KHS101 (7.5 μM, K) for 1 hour. Proteins within the highlighted KHS101-treated NP1 and GBM1 fractions were identified by mass spectrometry. (D) KHS101-GBM protein aggregation represented in a predicted (STRING) interaction network of proteins. Only GAPDH (italic) was shared between NP1 and GBM1 cells. Green, red, blue, and yellow colors represent enzymatic functions in protein folding, glycolysis, OXPHOS, and glycine metabolism, respectively. (E) qRT-PCR radar chart depicting KHS101-induced (7.5 μM) mRNA expression changes in GBM1 compared with NP1 cells. (F) Confocal analysis of tissue sections from vehicle-or KHS101-treated tumors (n=4, 3 sections per brain) indicates elevated DDIT3 immuno-positivity in tumor cells (P<0.01, Mann Whitney U-test, one tailed). Scale bar: 20 μm.

Furthermore, to confirm reduced HSPD1 function by KHS101 *in vivo*, we examined expression of DDIT3 (alias CHOP), a key marker of the unfolded mitochondrial protein response in glioma cells (26). Consistent with an increase of *DDIT3* expression in GBM cell culture (Fig. 4E), KHS101 induced a marked upregulation of DDIT3 protein levels in the GBM1 xenograft model (10) (Fig. 4F). In agreement with a causative relationship between chemical HSPD1 inhibition and bioenergetic failure in GBM cells, stable isotope substrate labeling with U-^13^C glucose (27) revealed that both glycolytic and TCA flux were selectively impaired in GBM1 cells 4 hours after KHS101 addition. Increased ^13^C enrichment of intracellular glucose was concomitant with decreased ^13^C incorporation into the glycolytic end-product lactate (Fig. 5). Moreover, the incorporation of the ^13^C-label into TCA cycle intermediates citrate and succinate markedly decreased in KHS101-treated GBM compared to NP cells (Fig. 5). This bioenergetic flux deficiency was accompanied by mitochondrial degradation (assessed by Immuno-EM 8 hours after KHS101 treatment; Supplementary Fig. 7A), and a reduction in GBM cell mitochondrial mass that was not evident in the NP control cells (assessed by silver staining 4 hours after KHS101 treatment; Supplementary Fig. 7B). Consistently, cellular ATP levels were markedly depleted in GBM1 compared to NP1 cells (Supplementary Fig. 7C).

**Fig. 5.**
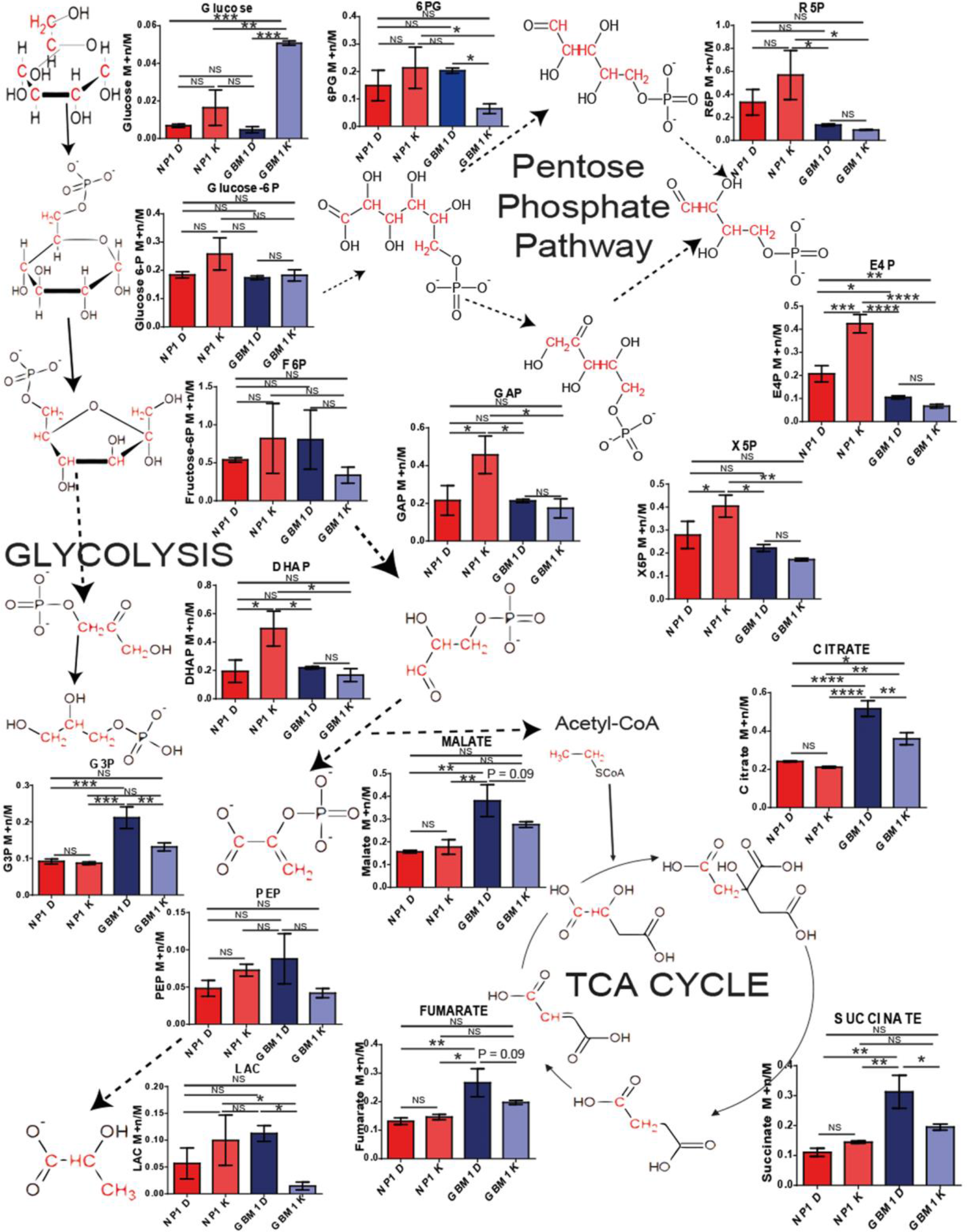
KHS101 specifically impairs glycolysis and the TCA cycle of GBM1 cells. Gas Chromatography-Mass Spectrometry stable isotope flux analysis of methoximation and silylation-derivatized metabolites extracted from NP1 (red bars) and GBM1 cells (blue bars) following a 4 hour treatment with or DMSO (D; 0.1%) KHS101 (K; 7.5 μM) in media containing U-13C glucose. KHS101 increases ^13^C enrichment of the intracellular glucose pool and decreases enrichment of glycerol-3-phosphate (G3P), the glycolytic end product lactate (LAC) and the TCA cycle intermediates citrate, succinate, fumarate and malate in GBM1 cells. Graphs show the M+n/M isotope ratio ^13^C enrichment of the following mass-to-charge (m/z) metabolite fragments: glucose 554 m/z C1-C6 M+6/M, glucose 6-phosphate (glucose 6P) 357 m/z C5-C6 M+2/M, fructose 6-phosphate (F6P) 459 m/z C4-C6 M+3/M, glyceraldehyde 3-phosphate (GAP) 400 m/z C1-C3 M+3/M, Dihydroxyacetone phosphate (DHAP) 400 m/z C1-C3 M+3/M, G3P 445 m/z C1-C3 M+3/M, phosphoenolpyruvate (PEP) 369 m/z C1-C3 M+3/M, LAC 219 m/z C1-C3 M+3/M, 6-phosphogluconate (6PG) 333 m/z C1-C4 M+4/M, Ribose 5-phosphate (R5P) 459 m/z C1-C3 M+3/M, Xylulose 5-phosphate (X5P) 604 m/z C1-C5 M+5/M, Erythrose 4-phosphate (E4P) 502 m/z C1-C4 M+4/M/. Data are mean ± SEM of 3 biological replicates. NS, not significant, *, P<0.05; **, P<0.01; ***, P<0.001; ****, P<0.0001, student's t-test (two tailed).

### KHS101 attenuates tumor proliferation and invasion *in vivo*

It has been shown that the mitochondrial unfolded protein response can be exploited for anti-glioma therapy (26). Accordingly, we investigated the pharmacological effects of KHS101 *in vivo* using xenograft tumors that were allowed to establish for 6 weeks after injection of GBM1 cells (1 × 10^5^) into the forebrain striatum. We adapted the KHS101 dosing regimen from previous neurogenesis work in rats (11) using a 10-week tumor treatment strategy (s.c., 6 mg/kg, b.i.d., and bi-weekly alteration of 5 and 3 treatment days per week). Immunohistological analysis of the vehicle-and KHS101-treated tumors at the 16-week endpoint showed that tumor cell proliferation was markedly reduced in KHS101-treated mice (^~^2-fold, Fig. 6A). This finding was consistent with a homogenous decrease in MKI67 expression and abrogation of clonal growth capacity in individually profiled GBM1 cells *in vitro* (Fig. 6B and C). KHS101-treated tumors showed signs of necrosis compared with tumors treated with vehicle control (indicated by a marked increase in nuclear-sparse areas; Fig. 6D, Supplementary Fig. 8A and B). The highly invasive phenotype of the GBM1 xenograft tumor model (10) enabled quantification of caudal tumor expansion and tumor cell migration across the corpus callosum into the contralateral hemisphere (a pathological hallmark of advanced GBM). KHS101 treatment markedly reduced both invasion of Vimentin-positive GBM1 cells into the corpus callosum of the contralateral hemisphere (≥2-fold; Fig. 6E) and caudal tumor expansion (Fig. 6F). Importantly, we did not observe signs of lethargy, weight loss, or other indicators of adverse effects over the treatment period and histological analysis demonstrated preserved hepatic architecture in KHS101-treated animals (Fig. 6G).

**Fig. 6.**
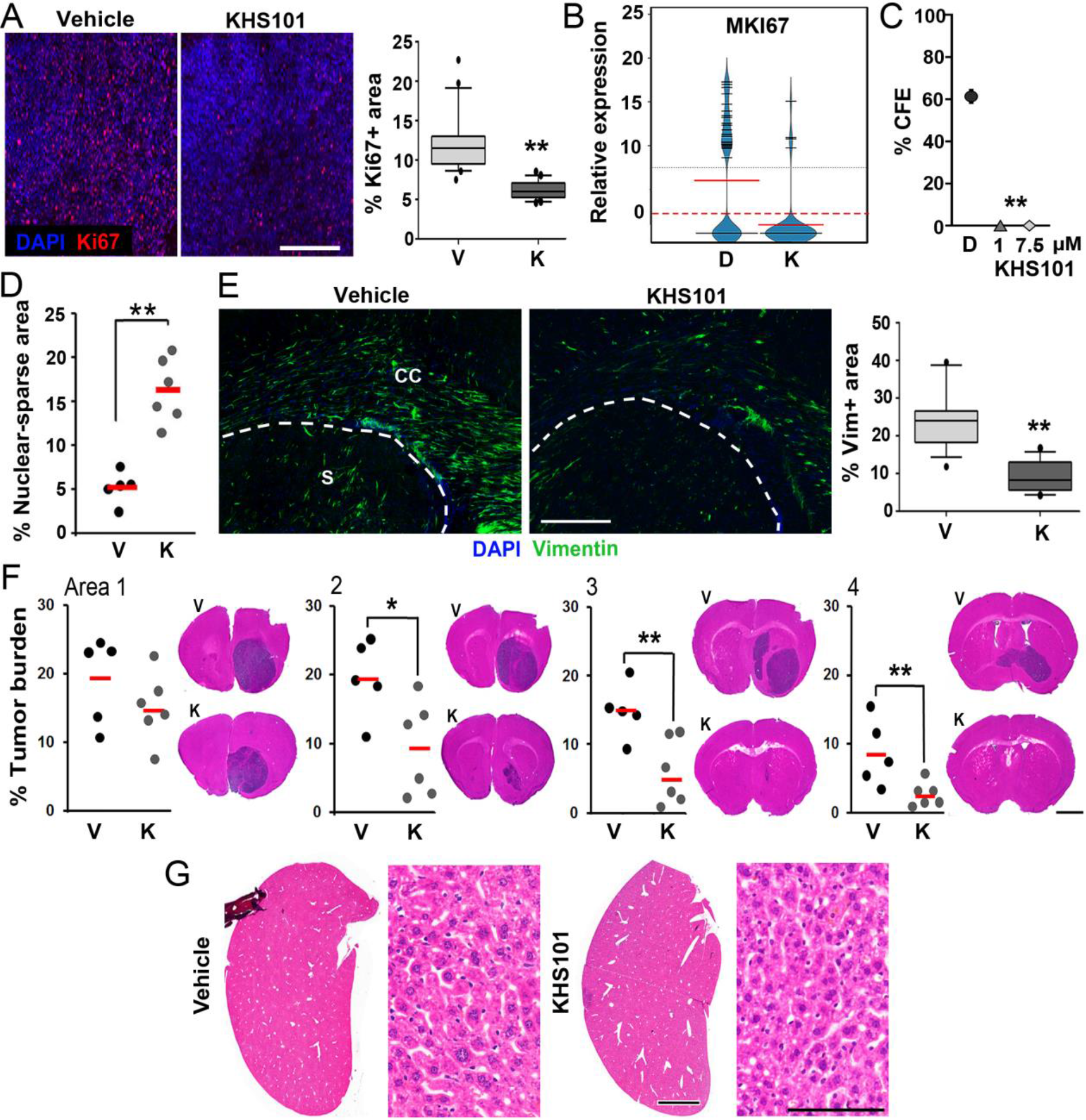
KHS101 significantly reduces GBM growth *in vitro* and *in vivo*. (A) Immunofluorescence images and quantification of vehicle-or KHS101-treated GBM1 tumors stained for Ki67. Scale bar, 400 μm. (B) Bean plot of *MKI67* single cell gene expression 5 days after KHS101 (7.5 μM) compared with DMSO (0.1%) treatment. (C) KHS101 treatment (1 or 7.5 μM) abrogates the clonal growth capacity of GBM1 cells compared to DMSO (0.1%)-treated cells. (D) Quantification of nuclear-sparse areas in anterior GBM1 tumor sections; V: vehicle, K: KHS101. Dots represent individual animals in control (black, n=5) and treatment (grey, n=6) groups. Red lines, mean. (E) Immunofluorescence images and quantification of Vimentin-positive GBM cells infiltrating the corpus callosum of the hemisphere contralateral to the injection site in KHS101-compared with vehicle-treated animals (n=4, 3 or 4 sections per brain). Dotted lines indicate border of corpus callosum (CC) and striatum (S). Scale bar, 300 μm. (F) Frontal to caudal GBM1 tumor burden in treated animals assessed by hematoxylin-eosin staining in 4 sequential brain areas; V: vehicle, K: KHS101. Dots represent individual animals in control (black, n=5) and treatment (grey, n=6) groups. Red lines, mean. Scale bar, 2 mm. (G) H&E staining of sections from vehicle-and KHS101-treated livers display unchanged hepatic architecture. Scale bars, 2 mm and 100 μm. Boxplots show the 10-90 percentile (black line, median), *, P<0.05; **, P<0.01, Mann Whitney U-test (one tailed).

Compared with normal brain, gliomagenesis is associated with elevated HSPD1 expression that negatively correlates with patient survival (Fig. 7A, Supplementary Fig. 8C). Accordingly, we further investigated whether KHS101 prolonged survival in a suitable xenograft model (characterized by a predictable onset of morbidity). To this end, we used a different (giant cell GBM-based) model, established with exclusively *in vivo*-propagated primary cells (GBMX1; onset of morbidity: 10-13 weeks). In two independent experiments (using early removal criteria within a moderate severity bandwidth), we found that the survival of animals carrying GBMX1-tumors (established 2 or 6 weeks before treatment) was markedly increased using the aforementioned treatment regimen for 10 weeks (Fig 7B), or continuously until the experimental endpoints (Fig. 7C). Histological endpoint analysis of KHS101-and vehicle-treated animals confirmed a significantly decreased tumor burden (^~^2-fold) in the animals that had received continuous KHS101 treatment (Fig. 7D). In summary, these results indicate significant anti-GBM effects of KHS101 *in vivo*, without discernible adverse toxicity.

**Fig. 7.**
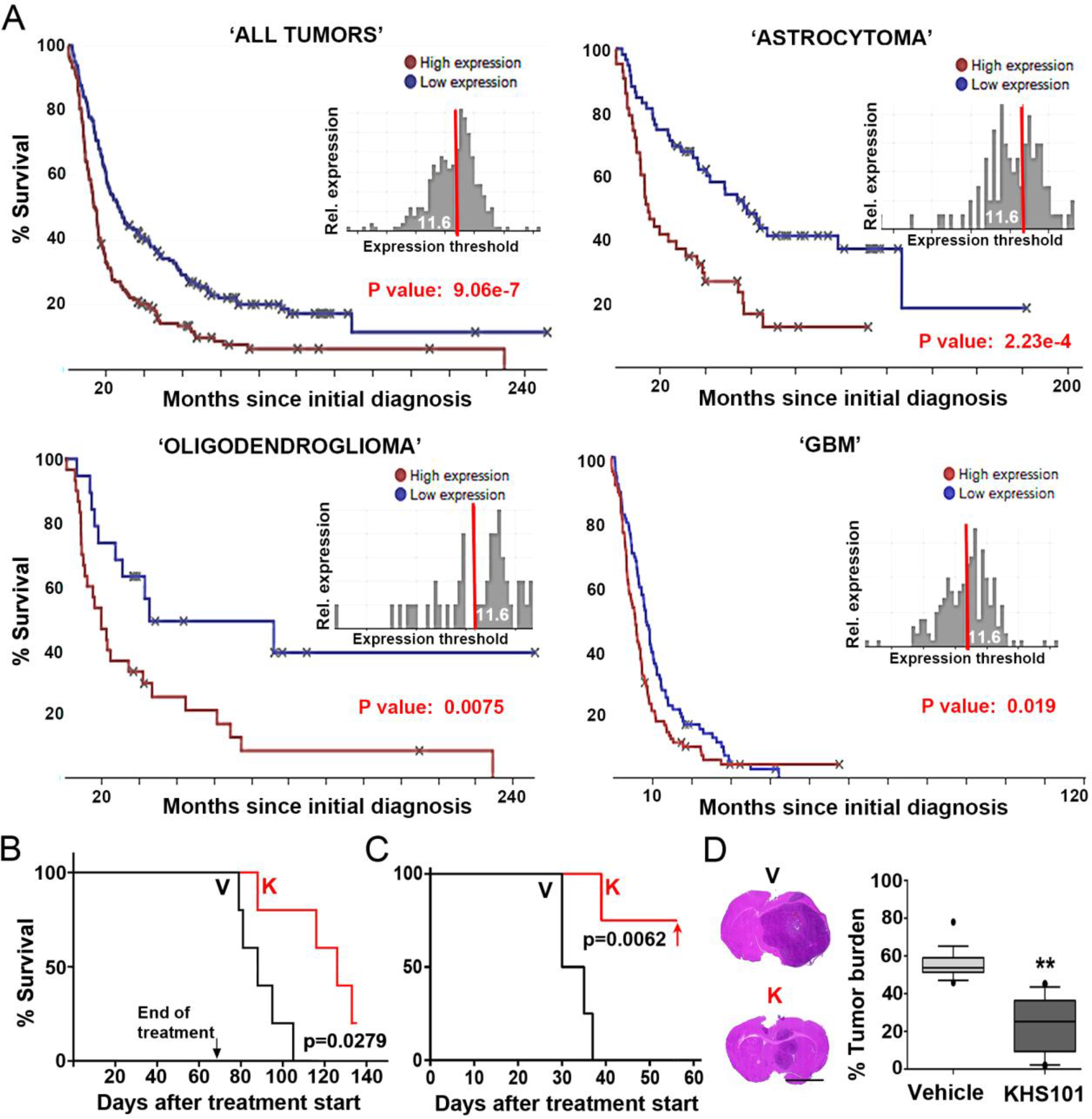
Elevated HSPD1 is associated with poor survival and KHS101 treatment improves outcomes in the GBMX1 model. (A) Kaplan Meier survival curves were retrieved from the ‘Project Betastasis’ web-portal (http://www.betastasis.com/glioma/rembrandt/kaplan-meier_survival_curve/) using the median preset threshold. Data shows that HSPD1 expression is negatively correlated with patient survival and the results are highly significant (p<0.01) for the ‘All tumor’, Astrocytoma, and Oligodendroglioma categories, and significant (p≤0.02) for GBM tumors. (B) Kaplan-Meier survival curves of GBMX1 tumor-carrying animals. Tumors were established over a 2 week period followed by 10 weeks KHS101 (K) or vehicle (V) treatment. (n=7 animals per group; P=0.0279, log-rank test). (C) GBMX1 tumors were established over a 6 week period followed by continuous KHS101 (K) or vehicle (V) treatment until the experimental endpoint (indicated by the red arrow). The log-rank test was used to calculate the p value. (D) H&E stained sections (n=4, 4 or 5 sections per brain) and quantification of tumor burden in corresponding animals. Scale bar, 3 mm. Boxplot shows the 10-90 percentile (black line, median), *, P<0.05; **, P<0.01, Mann Whitney U-test (one tailed).

## Discussion

GBM is a devastating cancer with limited treatment options and correspondingly poor patient outcomes. We based our investigation on the hypothesis that a synthetic small molecule can significantly suppress GBM tumor progression and focused on the compound KHS101 that shows BBB penetrability as well as non-toxic neuronal differentiation properties *in vitro* and *in vivo* (11). One challenge for GBM target discovery and validation is to incorporate the ever-changing composition of molecularly and phenotypically diverse tumor cell populations (12, 13) into preclinical disease modelling. However, transcriptional diversity among and within our six patient-derived models was revealed by microfluidic single cell qRT-PCR analysis. We developed computational approaches that robustly indicated classical, proneural, and mesenchymal GBM subtype compartments in our GBM models (Supplementary Report 1). Notably, KHS101 cytotoxicity was independent of GBM subtype features, parental tumor origin (primary versus recurrent GBM), and MGMT methylation status (Fig. 1). Neither pro-differentiation signaling (10, 15) nor low oxygen culture conditions (that can promote GBM clonogenicity (28)) significantly changed the GBM cell sensitivity to KHS101. Overall, these results point towards a molecular mechanism that is distinct from ‘forced’ GBM stem cell-like differentiation (a non-cytotoxic phenotype that develops over several days (10)). Consistently, the KHS101-mediated downregulation of the neural stem cell-promoting factor TACC3 over 48 hours was not involved in the immediate metabolic response characterized by mitochondrial dysfunction and bioenergetic deficiency in GBM compared with NP cells.

In agreement with KHS101-induced mitochondrial stress, affinity-based target identification revealed a physical interaction between KHS101 and the mitochondrial chaperone HSPD1 (Fig. 2). In line with this finding, KHS101 rapidly inhibited enzymatic HSPD1 activity without affecting the chaperone’s protein or gene expression levels. These results highlight the significance of recognizing and exploiting differences between genetic and small-molecule target inhibition (e.g., reviewed in (29, 30)). Accordingly, the HSPD1-targeting properties of KHS101 were further strengthened by an independent chemical positive control (MC) that recapitulated the KHS101-induced GBM cell degradation phenotype. Intriguingly, MC (a non-prenylated acylphloroglucinol) and KHS101 (a thiazole derivative) share anti-cancer and mitochondrial HSPD1-targeting properties (21, 31), and it is unlikely that these chemically-distinct entities phenocopy each other due to ‘off-target’ effects (29). In agreement with the observed concentration-dependent inhibition of HSPD1 chaperone activity by KHS101, we detected a significant selective aggregation of mitochondrial proteins in GBM cells 1 hour after KHS101 addition. The attenuation of activities facilitated by enzymes such as ALDOA (regulating glycolysis), DLST (regulating TCA cycle), SLC25A3 and ATP5A1 (regulating OXPHOS), and the chaperone LONP1 (regulating mitochondrial integrity in cancer cells) were consistent with the wide-ranging metabolic stress observed in the GBM but not NP cells (Fig. 3). In addition, this GBM mitochondrial unfolded protein response was in line with the observed KHS101-induced upregulation of DDIT3 (alias CHOP) expression *in vivo*, corroborating evidence for a link between mitochondrial homeostasis and GBM cellular fitness (26, 32). Importantly, selective effects of KHS101 towards brain cancer cells were observed throughout our study at protein, metabolite, mRNA, and organelle level. In agreement with these findings, KHS101 is not toxic in other non-cancer contexts (11, 17). For example, previous studies afforded to KHS101 showed favorable *in vivo* properties including accelerated neuronal differentiation in adult rats (without affecting apoptosis of brain cells; (11)). Consistent with the specific KHS101 cytotoxicity in GBM compared with NP cells, the compound markedly decreased the progression of established xenograft tumors and adverse effects (e.g., liver toxicity) were not observed in treated mice after prolonged (10-week) administration (Fig. 4).

In summary, this experimental small molecule phenotype and target profiling study identifies HSPD1 enzymatic function as a specific molecular vulnerability in GBM cells. The data indicate that a lethal GBM cell fate can be selectively triggered in a heterogeneous spectrum of GBM cells by a single agent. The cytotoxic activity of KHS101 was selective for tumor cells highlighting a dependency on mitochondrial HSPD1 chaperone activity that critically facilitates GBM cell energy metabolism. These findings may be exploited for therapeutic development(s), and future studies are warranted to address the role of HSPD1 in the metabolic reprogramming driving brain tumorigenesis.

## Methods

### Patient-derived GBM cell models

Adherent culture of the GBM1, GBM4, and the NP cell models has been previously described (14). Additonal models were derived under the governance of the ethically-approved Leeds Multi-disciplinary Research Tissue Bank using similar methodology. U-87 MG (ECACC, 89081402) and U251 cells were cultured in DMEM medium supplemented with 10% (v/v) fetal bovine serum. All cell lines were confirmed as mycoplasma negative.

### Single cell isolation and profiling

Individual GBM cells were captured from cell suspensions using the microfluidic Fluidigm C1 single-cell auto prep system (Fluidigm). Reverse transcription and pre-amplification were carried out within a 96-well microfluidic C1 chip according to the manufacturer’s instructions using DELTAgene assays (Fluidigm, Supplementary Table 2).

Reverse transcription and pre-amplification were carried out within a 96-well microfluidic C1 chip (10-17 μM) according to the manufacturer’s instructions. Pre-amplified cDNA samples from single cells were analyzed by qRT-PCR using 96.96 Dynamic Array™ IFCs and the BioMark™ HD System (Fluidigm). Each analysis comprised up to 96 cDNA samples from individual cells and DELTAgene assays (Supplementary Table 2). Amplified cDNA (3.3 μL) was mixed with 2x Ssofast EvaGreen Supermix (2.5 μL), Low ROX buffer (2.5 μL; Bio-Rad, PN 172-5211, Hercules, CA, USA), and ‘sample loading agent’ (0.25 μL; Fluidigm, PN 100–3738). DELTAgene forward and reverse primers were mixed with Fluidigm Assay Loading Reagent (2.5 μL). Samples and assays were loaded onto Fluidigm M96 chips using the HX IFC Controller (Fluidigm) and then transferred to the BioMark™ HD real-time PCR reader following the manufacturer’s instructions. PCR was performed using the (GE Fast 96 × 96 PCR + Melt v2.pcl) thermal protocol: Thermal Mix of 70°C, 40 minutes; 60°C, 30 seconds, Hot Start at 95°C, 1 minute, 30 cycles of (96°C, 5 seconds; 60°C, 20 seconds), and ‘Melting’ using a ramp from 60°C to 95°C at 1°C/3 seconds. Data were analyzed using the Fluidigm Real-Time PCR Analysis software and Ct and melting-curve data were exported to Excel and R for further analysis. Results were assessed using the Fluidigm Real-Time PCR Analysis software (heat map view). Melting curve analyses was carried out to identify non-specific amplicons, which were removed from the final data set. For calculation of relative expression values (ΔCt = LoD Ct - Ct), the limit of detection (LoD) was set to a Ct value of 30 (Ct values of ‘999’ were allocated artificial values ≥30 to allow for the visualization of ‘non-expressors’ in bean plots). Plots were generated using BoxPlotR (33)

### Computational single cell gene expression analysis

After transformation of Ct to expression values, missing values (e.g., resulting from unspecific amplicons removed after melting curve analysis) were imputed using k-nearest neighbor imputation (34). Single-cell expression levels were adjusted for cell cycle-dependent heterogeneity as described in (35) using 20 known cell cycle markers. Gene expression levels of tumor samples with subtype annotation were obtained from (14) and integrated with our single cell qPCR data set. To this end, both data sets were separately discretized to three levels on a per gene basis using mixture models. Subsequently, a Random Forest classifier was trained on the data set reported by Verhaak and colleagues (14), and used to predict the subtypes of the single cells. Variability of gene expression was analyzed by (i) calculating the fraction of cells expressing a certain gene among all cells with a non-missing measurement, and (ii) calculating the coefficient of variation among all cells with a non-zero expression value. Complete code and data for our computational approaches are shown in Supplementary Report 1.

### Electron Microscopy

For standard EM cell pellets were fixed in 4% (w/v) formaldehyde/glutaraldehyde solution in 100 mM phosphate buffer at room temperature (RT) for 30 minutes, and subsequently washed (2 × 10 minutes) and stored at 4°C in 100 mM phosphate buffer. Cells were then post-fixed with 1% buffered osmium tetroxide on ice for 45 minutes, washed in 100 mM phosphate buffer (2 × 10 minutes) then treated with buffered 1% tannic acid solution (10 minutes) and washed again (100 mM phosphate buffer; 2 × 10 minutes). Cell samples were stained with 2% uranyl acetate and dehydrated in increasing concentrations of ethanol/acetone before embedding. These samples were then polymerized overnight at 70°C. Processed cell pellets were sectioned to 1 μm on a Leica EM UC7 microtome and stained with toluidine blue. Further thin sections were then taken (70 nm) and placed on 300 mesh uncoated copper grids, and these were stained with saturated uranyl acetate in 50% ethanol followed by Reynolds lead citrate for 10 mins each. For Immuno-EM cell pellets were fixed in 4% (w/v) formaldehyde/ glutaraldehyde solution in 100 mM phosphate buffer on ice then stored at 4°C. Cells were then washed and resuspended in 12% gelatin for 15 minutes at 37°C, spun down, cut and left in 2.3 M sucrose overnight at 4°C. Cell blocks were mounted on Leica holders and frozen in liquid nitrogen. Sections (95 nm) were cut at −90°C prior to immunolabeling with anti-COX IV primary antibody (Cell Signaling; 4850; 1:50), and subsequently gold labeled anti-rabbit secondary antibody (BBI solutions; EM.GAR10; 1:20). Sections were post-fixed in 1% (v/v) glutaraldehyde solution in sodium phosphate buffer for 10 minutes, washed 3 times in dH_2_O then further in 2% methyl cellulose in 0.4% uranyl acetate (10 minutes). All sections were imaged using a FEI Tecnai 12 transmission electron microscope at an accelerating voltage of 120 kV.

### Cytogenetic analysis of tumor cells

GBM cells and NP1 cells were analyzed for alterations in chromosomes 7 and 10 using fluorescence in-situ hybridization (FISH) methodologies for EGFR and PTEN respectively. Cells were grown to 50% confluency and then exposed to 0.2 mg/ml colcemid for four hours at 37°C. Subsequently, cells were collected following 3 minutes exposure to trypsin/EDTA (Gibco; 15400054). Resultant cell suspensions were harvested using 15 minutes exposure to a 50:50 mixture of pre-warmed 0.075 M KCl/1% Sodium Citrate, followed by 3 washes in Carnoy’s fixative. Slides previously stored in absolute ethanol were used to make preparations from the fixed cell suspensions (60°C for 30 minutes). EGFR FISH studies were carried out using the directly labelled Kreatech EGFR (7p11) / SE7 probe. PTEN FISH studies were carried out using the Vysis PTEN / CEP10 FISH Probe Kit according to manufacturer’s instructions. Interphase cells (^~^100) were analyzed by epifluorescence microscopy for signal imbalance. Deletions were called when seen in over 20% of cells. Amplification was defined as over four times the copy number for a given cell ploidy.

### Chemical synthesis

All chemical reagents for synthesis were purchased from commercial suppliers and used without further purification. When used as reaction solvents, THF and DCM were dried and deoxygenated using an Innovative Technology Inc. PureSolv^®^ solvent purification system. Flash column chromatography was carried out using silica (Merck Geduran silica gel, 35–70 μm particles). Thin layer chromatography was carried out on commercially available pre-coated aluminium plates (Merck silica 2 8 8 0 Kieselgel 60 F_254_). Lyophilisation of compounds was performed using a Virtis Benchtop K freeze dryer. Automated RP (C18) chromatography was performed using a Biotage Isolera™ Prime Advanced Flash Purification system with RediSep Rf C_18_ columns (26 g or 130 g; Teledyne Isco). Compounds were eluted with the indicated solvent mixtures/gradients; elution was monitored by UV detection and fractions were analysed by LC-ESI-MS before combination and concentration of fractions containing pure product.

Preparative high performance liquid chromatography (HPLC) was performed on an Agilent 1100 Infinity Series equipped with a UV detector and Ascentis Express C18 reverse phase column using MeCN/water (5→95%) containing 0.1% formic acid. A flow rate of 0.5 mL min^−1^ was used over a period of 8 minutes. Analytical HPLC was performed on an Agilent 1290 Infinity Series equipped with a UV detector and a Hyperclone C18 reverse phase column using MeCN/water (5→95%) containing 0.1% formic acid, at either 0.5 mL min^−1^ over a period of five minutes or 1.0 mL min^−1^ over a period of 30 minutes.

High resolution electrospray (ESI+) mass spectrometry was performed on a Bruker MaXis Impact QqTOF mass spectrometer, and *m/z* values are reported in Daltons to four decimal places. LC-ESI-MS data were recorded on an Agilent Technologies 1200 series HPLC combined with a Bruker HCT Ultra ion trap using 50 × 20 mm C_18_ reverse phase columns using MeCN/water (5→95%) containing 0.1% formic acid. A flow rate of 1.5 mL min^−1^ was used and *m/z* values are given in Daltons to one decimal place.

Microwave reactions were performed using a Discovery Explorer 48 Autosampler (CEM Corporation) single-mode microwave instrument producing controlled irradiation at 2450 MHz, using standard sealed microwave glass vessels. The instrument was operated in temperature control mode and reaction temperatures were monitored with an IR sensor on the outside wall of the reaction vessel. Reaction times refer to hold times at the indicated temperatures, not to total irradiation times.

^1^H and ^13^C NMR spectra were recorded in deuterated solvents on a Bruker Avance 500 or Bruker Avance DPX 300. Chemical shifts are quoted in parts per million downfield of tetramethylsilane and referenced to residual solvent peaks (CDCl_3_: ^1^H = 7.26 ppm, ^13^C = 77.16 ppm, DMSO-d_6_: ^1^H = 2.50 ppm, ^13^C = 39.52 ppm) and coupling constants (*J*) are reported to the nearest 0.1 Hz. Assignment of spectra was based on expected chemical shifts and coupling constants, aided by COSY, HSQC and HMBC measurements where appropriate.

The synthesis of KHS10, KHS101-BP, and KHS101-biotin is described in the Supplementary Information.

### Cellular assays

For live cell analysis, cells were allowed to grow for 2 days before the addition of KHS101 (7.5 μM) or DMSO (0.1%), and subsequently monitored for 3 days. Images were acquired at 45 minute intervals using the IncuCyte ZOOM^®^ live cell imaging system (Essen Bioscience). Cellular growth rates per hour (GR, %) were calculated for a 60 hour period: GR = [78-hour-confluency (%) – 18-hour-confluency (%)] / [18-hour-confluency (%) × 60 hours]. Confluency curves were based on relative values (normalized to the t_0_ time point).

For the analysis of cell viability and caspase 3/7 activation, cells were seeded into 96 well plates (Greiner bio-one; 655073) at densities of 10,000 and 2,500 cells, respectively. The following day, cells were treated with vehicle (DMSO) or KHS101 (1-20 μM) in 100 μL of medium and the CellTiter-Glo^®^ and Caspase-Glo^®^ 3/7 assays (Promega) were carried out at the indicated time points according to the manufacturer’s instructions.

For the quantification of apoptosis using annexin V and propidium iodide, GBM1 cells were treated with KHS101 (7.5 μM) or vehicle (DMSO, 0.1%) for 48 hours, then harvested with trypsin, washed with PBS, and stained with Annexin V and Propidium Iodide for 15 minutes at 37°C using an Annexin V-fluorescein staining kit (Roche) in accordance with the manufacturer’s protocol. Labelled early apoptotic and late apoptotic/necrotic cells were quantified through quadrant gating using a NC3000 cytometer (Chemometec).

Autophagic kinetics were assessed using the CytoID^®^ detection assay (Enzo Life Sciences) according to the manufacturer’s instructions (1:500 dilution). Stained live cells were imaged using an EVOS digital inverted fluorescence microscope.

For the differentiation assay, cells were seeded into 96 well plates (Greiner bio-one; 655073) at a density of 5,000 cells per well and treated with recombinant human BMP4 at 100 ng/mL for 4 days. Subsequently, cells were treated for 48 hours with DMSO (0.1%) or KHS101 (1-20 μM) in 100 μL of medium and the CellTiter-Glo^®^ assay (Promega) was carried out according to the manufacturer’s instructions.

For the colony formation assay, cells were seeded at a density of 125 cells/well in 24-well plates and allowed to adhere. The following day, the single cells per well were counted and treated with DMSO or KHS101. Colonies consisting of >6 cells were counted after 10 days and the percentage of cells that were able to form a colony was determined.

### Gene Expression analysis

Gene expression profiling was carried out using the Illumina HumanHT-12 v4 beadchip. Raw data were pre-processed with a quantile normalization method using Bioconductor. Differential analysis log fold changes were calculated for all probes and those with a false discovery rate (FDR) ≤ 0.075 were selected.

For qRT-PCR analysis, total RNA was extracted using the RNeasy mini kit (Qiagen) according to the manufacturer’s instructions. Copy DNA was synthesized using the SuperScript II first-strand-synthesis method with oligo(dT)s (Invitrogen) and analyzed using TaqMan^®^ gene expression assays (Supplementary Table 3). High-throughput qRT-PCR was carried out using the Fluidigm BioMark™ HD System. Data were internally normalized to Glyceraldehyde-3-phosphate dehydrogenase (GAPDH).

### Immunocytochemistry

After fixation with 4% (w/v) paraformaldehyde (10 minutes, RT), cells were permeabilized with ice-cold methanol (10 minutes; −20°C) for LC3B (Cell Signaling; 2775S; 1:200). Non-specific antibody binding was reduced in PBS ‘blocking buffer’ containing 10% (v/v) FBS and 0.03% (v/v) Triton X-100 at RT for 1 hour. Primary antibody was used in ‘blocking buffer’ at 4°C overnight. Secondary antibody: Cy3-conjugated (Jackson ImmunoResearch; 711-165-152; 1:400). Nuceli were stained using DAPI (Sigma; 1 μg/mL).

Images were acquired using an EVOS digital inverted fluorescence microscope (life technologies) or a Nikon A1R confocal microscope.

### TACC3 and HSPD1 lentiviral-based RNAi

Generation of lentiviral particles was carried out according to the manufacturer’s specifications and as previously described (10). After 16 hours, medium containing residual virus was washed out and replaced with fresh media containing 2 μg/mL puromycin. Protein and mRNA knockdown efficiencies were determined by Western blotting and qRT-PCR at the indicated time points.

*The two different lentiviral TACC3 small hairpin (sh) RNA and non-targeting constructs were*:

shTACC3#1 [TRCN0000062026, Sigma]
Sequence: CCGGGCTTGTGGAGTTCGATTTCTTCTCGAGAAGAAATCGAACTCCACAAGCTTTTTG
shTACC3#2 [TRCN00000 62024, Sigma]
Sequence: CCGGGCAGTCCTTATACCTCAAGTTCTCGAGAACTTGAGGTATAAGGACTGCTTTTTG
shSCR [SHC016, Sigma]
Sequence: CCGGGCGCGATAGCGCTAATAATTTCTCGAGAAATTATTAGCGCTATCGCGCTTTTT

*The four different lentiviral HSPD1 shRNA and non-targeting constructs were*:

shHSPD1#1 [TRCN0000343883, Sigma]
Sequence: CCGGCCTGCTCTTGAAATTGCCAATCTCGAGATTGGCAATTTCAAGAGCAGGTTTTTG
shHSPD1#2 [TRCN0000343882, Sigma]
Sequence: CCGGGCTAAACTTGTTCAAGATGTTCTCGAGAACATCTTGAACAAGTTTAGCTTTTTG
shHSPD1#3 [TRCN0000343951, Sigma]
Sequence: CCGGCCAGCTTAAAGATATGGCTATCTCGAGATAGCCATATCTTTAAGCTGGTTTTTG
shHSPD1#4 [TRCN0000343952, Sigma]
Sequence: CCGGGCAATGACCATTGCTAAGAATCTCGAGATTCTTAGCAATGGTCATTGCTTTTTG
shSCR [SHC016, Sigma]
Sequence: CCGGGCGCGATAGCGCTAATAATTTCTCGAGAAATTATTAGCGCTATCGCGCTTTTT

### Western blotting

Complete cellular extracts (^~^10^6^ cells) were lysed in CellLyticMT (Sigma; C3228) plus protease/phosphatase inhibitor cocktail (Thermo Scientific; 78440) following the manufacturer’s recommendations. To achieve nuclear/cytoplasmic separation of total cellular protein, the CelLytic ™ NuCLEAR ™ Extraction Kit (Sigma; NXTRACT-1KT) was used. For mitochondrial/cytosolic fractionation the Mitochondria Isolation Kit for Cultured Cells (Thermo Scientific; 89874) was utilized. Lysate concentrations were determined using the BCA protein assay (Pierce; 23225). Protein (between 15-30 μg) was loaded onto Mini-Protean TGXTM pre-cast gels (10%; Biorad), and transferred onto a nitrocellulose membrane (0.45 μm, Biorad). Membranes were exposed to the following antibodies: mouse anti-TACC3 (Santa Cruz; sc-48368; 1:500), mouse anti-Actin beta (Sigma; A5441; 1:20,000), rabbit anti-COX IV (Cell Signaling; 4850; 1:500), rabbit anti-Hsp60 (Abcam; ab46798; 1:500) combined with secondary anti-rabbit (Thermo; G-21234; 1:2000) or anti-mouse (Sigma; A4416; 1:5000) antibodies conjugated to horseradish peroxidase. Membranes were developed using the Luminata Crescendo or Forte Western HRP substrate solutions (Millipore).

### Quantification of metabolic phenotypes

Metabolic analysis in real-time was carried out using the XFp extracellular flux analyser (Seahorse Bioscience). Changes to mitochondrial function or glycolytic flux where assessed with ‘Mito Stress Test’ or ‘Glycolysis Stress Test’ kit protocols (Seahorse Bioscience 101848-400 and 103020-400) using seeding densities of ^~^30,000 cells per well. Prior to analysis, the culture medium was replaced with XF base media (Agilent; 102353-100) and the cells transferred to a 37°C non-CO_2_ humidified incubator. For the ‘Mito Stress Test’ XF base media was supplemented with glucose (25 mM; Sigma; G8769) and sodium pyruvate (0.5 mM; Sigma; S8636). L-glutamine (2 mM; Gibco; 35050-061) was used for both ‘Mito and Glycolysis Stress Tests’ with NP cells. Supplemented culture media were adjusted to pH 7.4 and filtered through a 0.2 μm filter, and maintained at 37°C throughout the experiments. KHS101 and vehicle (DMSO) were injected at the indicated concentrations after the baseline metabolic flux readings were established. Oligomycin (1 μM), FCCP (0.5 μM), antimycin and rotenone (0.5 μM) were injected according to the ‘Mito Stress Test’ protocol. Glucose (10 mM), oligomycin (1 μM) and 2-deoxy-glucose (100 mM) were sequentially injected according to ‘Glycolysis Stress Test’ protocols.

Gas Chromatography-Mass Spectrometry stable isotope flux analysis of methoximation and silylation-derivatized metabolites was carried out as previously described (27). Metabolites were extracted from NP1 and GBM1 cells following a 4 hour treatment with KHS101 (7.5 μM) or DMSO (0.1%) in media containing U-^13^C glucose.

Cellular ATP was determined 24 hours after treatments with KHS101 using a luminescence-based readout as described above (CellTiter-Glo^®^ assay; Promega). Data were normalized to DMSO controls.

### Thermal stability analysis

The NanoTemper Technologies’ Prometheus NT.48 instrument was used for studying the compound’s effect on HSPD1 thermal stability. KHS101 or vehicle (DMSO) was added to 30 μl of a 1 mg/mL HSPD1 solution at the indicated concentrations. The thermal stability of HSPD1 was observed in a 1°C/min thermal ramp from 20°C to 95°C with an excitation power of 75%. The emission wavelengths of tryptophan fluorescence at 330 nm and 350 nm were measured. Tm values were calculated from first derivative automatically using the PR.Control software.

### Histology and immunohistochemistry

After fixation with 4% (w/v) PFA at 4°C for two days, brains were placed in a sucrose solution (25% sucrose in 0.5 M NaH_2_PO_4_, 0.5 M Na_2_HPO_4_) until they had sunken to the bottom of the tube. Subsequently, the brains were snap frozen and cut as 30 μm sections on a cryostat. The sections were stored at −20°C in Walter’s antifreeze solution (13 mM NaH2PO4, 55 mM Na_2_HPO_4_, 30% (v/v) ethyleneglycol, 30% (v/v) glycerol). Prior to staining, the sections were equilibrated in PBS for 10 minutes. Sections were either stained with Hematoxylin and Eosin using standard protocols or subjected to immunohistochemical staining. For immunohistochemistry, tissue sections were permeabilized with PBS containing 0.2% (v/v) Triton X-100 for 10 minutes, followed by blocking with PBS containing 10% (v/v) FBS and 0.03% (v/v) Triton X-100 at RT for 1 hour. Incubation with primary antibodies in PBS containing 10% (v/v) FBS and 0.03% (v/v) Triton X-100 was carried out at 4°C overnight. The following antibodies were used: anti-Ki67 (Abcam; ab16667; 1:200), anti-Vimentin (DAKO; M0725; 1:200) and anti-DDIT3 (Cell Signaling; 2895; 1:1000). Secondary antibodies used were AlexaFluor488-conjugated (Molecular Probes; A11029; 1:200), AlexaFluor647-conjugated (Molecular Probes; A21235; 1:200) or Cy3-conjugated (Jackson ImmunoResearch; 711-165-152; 1:400). Nuclei were stained using DAPI (Sigma; 1 μg/mL) and whole sections were analyzed using the EVOS digital inverted fluorescence microscope or Nikon A1R confocal microscope and selected slides were scanned using the Leica Ariol system at a magnification of x20 with a resolution of ~0.35 microns per pixel.

### Image analysis

Immunocytochemistry: Regions of interest (ROIs such as LC3B positive area) were manually defined or by color thresholding using ImageJ (default settings, color space: HSB), and their percentage of the total cellular area was calculated. The number of MKI67 or CytoID^®^ positive cells was determined in randomly selected images totalling ≥500 cells per single experiment.

Immunohistochemistry: ROIs (KI67 and Vimentin-stained areas, eosin-dense regions, DDIT3-positive areas) were isolated from immunofluorescence/phase contrast or immunohistological images. The investigators were blind to the identity of the images which were analyzed using ImageJ software and color thresholding with default settings (color space: HSB). Selected ROIs were measured and the percentage of the total area of interest was determined. Five different tumor sections per specimen were analyzed for the MKI67-positive ROI. For measuring DDIT3-positive areas, 3 different tumor sections (each containing ^~^4 confocal imaging fields) were analyzed per specimen. For determination of GBM cell invasion in xenograft tumors, a minimum of 3 Vimentin-stained sections per specimen were analyzed.

### Affinity-based target identification

GBM1 cells were incubated with KHS101-BP (5 μM) in the presence or absence of unlabeled KHS101 (250 μM) for 30 minutes, and irradiated with UV light (365nm) for 30 minutes. Cells were lysed using 0.5% Triton X-100 and protease inhibitor cocktail (Sigma, cat # P8340). Cell lysates were incubated with 25 μM biotin azide (ThermoFisher, cat # B10184), 1 mM TCEP, 100 mM ligand (TBTA), and 1 mM aqueous copper sulfate at 4°C overnight. Subsequently, proteins were fractionated using ammonium sulfate and the 20-40% fractions were subject to 2D SDS/PAGE. Biotin-labeled proteins were detected through Western blotting using Abcam ab1227). Protein spots corresponding to the specific biotin-labeled proteins were visualized with silver staining on parallel gels. A distinct spot was excised and protein identified using liquid chromatography tandem mass spectrometry.

### Human HSPD1 and KHS101 *in vitro* binding

A total of 1 μg recombinant HSPD1 (Abcam; ab113192) was diluted in 1 mL PBS (with 2mM MgCl2, 2 mM DDT, and 0.1% tween 20) and incubated with 5 μM biotinylated KHS101 at 4°C overnight in the presence or the absence of non-labelled KHS101. Streptavidin agarose beads were added to the incubation mixture and rotated at 4°C for 2 hours. The beads were then precipitated and washed three times in PBS. Bound proteins were eluted with 2x SDS sample buffer and analyzed with SDS/PAGE followed by silver staining and Western blotting.

### HSPD1 protein refolding assay

The effect of KHS101 on HSPD1 mediated protein folding was assessed using the human HSP60/HSP10 protein refolding kit (R&D Systems, Cat # K-300) following the manufacturer’s instructions (no unspecific effect of KHS101 on the luminescence signal of client protein in absence of heat-induced protein denaturation was observed).

### Mitochondrial protein fractionation

Mitochondrial organelles were isolated from 10^7^ cells treated for 1 hour with DMSO (0.1%) or KHS101 (7.5 μM) using the Thermo Scientific Mitochondrial Isolation Kit (#89874). Mitochondrial preparations were normalized to the protein content and detergent soluble and insoluble fractions were separated by NP40 (v/v) treatment at indicated concentrations. Incubation was on ice for 20 min, and centrifugation at 12000 x g for 20 minutes at 4°C. Supernatant and pellet were separated and analysed by SDS-PAGE electrophoresis and silver staining.

### Xenograft tumor experiments

Animal experiments were carried out under UK project license approval and institutional guidelines. Animals were maintained under standard conditions (12 hour day/night cycle with food and water ad libitum). Experiments were carried out using 6 to 8-week-old NOD scid gamma (NSG) and BALB/c Nude mice for the GBM1 and GBMX1 models, respectively. Mice were stereotactically injected with 2 × 10^5^ GBM1 cells or 8 × 10^4^ GBMX1 cells in a volume of 2 μL (containing 30% Matrigel™; BD Biosciences) into the right striatum (2.5 mm from the midline, 2.5 mm anterior from bregma, 3 mm deep). Surgery was performed under general anaesthesia using aseptic techniques. Mice were monitored daily for signs of sickness, pain or weight loss. After the indicated tumor-establishing period, 6 mg/kg KHS101 or vehicle control (5% (v/v) ethanol, 15% (w/v) (2-Hydroxypropyl)-β-cyclo-dextrin) was administered subcutaneously (s.c.) twice daily with bi-weekly alteration of 5 and 3 treatment days per week. Experiments were concluded at indicated endpoints and tissue was subjected to immunohistological and image analysis.

### Statistical analysis

For *in vitro* data, a minimum of 3 independent biological repeats were analyzed using the student’s t-test (two tailed, equal variance) and expressed as mean ± SD. One biological repeat comprised a minimum of 3 technical replicates. Approximate normal distribution of data was assumed, however for small sample sizes (n<5), individual data points are identified in the figures. For xenograft tumor analysis, the Mann-Whitney U-test was used (one tailed). For Kaplan-Meier xenograft tumor analysis, the significance was calculated using the log-rank test. P values of ≤ 0.05 were considered significant (*) and P values of ≤ 0.01 were considered highly significant (**).

## Author’s contributions

E.S.P., V.B.K., C.A., R.K.M., H.A.B., E.C., J.W., B. DS., A.P., A.J. D., H.B.S.G., M.L., H.S., L.D.R., R.S.B., S.J.A., S.Z., F.M., and H.W. performed and/or coordinated the experimental work. E.S.P., V.B.K., C.A., E.M.R., E.C., H.S., L.D.R., R.S.B., S.J.A., S.Z., F.M., and H.W. carried out data analysis. E.M.R. and F.M. generated computer code. P.C., A.D., S.C.S. and HW collected and managed data, and provided materials. E.S.P., V.B.K, R.K.M., and H.W. prepared the manuscript and figures. R.S.B., J.G., S.J.A., S.Z., F.M., and H.W. provided project leadership.

## Acknowledgements

We thank J. Jauch for kindly providing myrtucommulone, S. Wilkinson for help with LC3/autophagy protocols, D. J. Beech for microscopy support and resources, P. Roberts and K. Rankeillor for help with cytogenetic analysis, P. O’Toole and M. Stark for help with EM imaging, H. Payne for help with qRT-PCR protocols, C. Simmons for assistance with chemical synthesis, and N. Riobo-Del Galdo, P. Ceppi, and P.J. Selby for useful discussions. H.W. acknowledges support from the MRC New Investigator Award (MR/J001171/1), the Marie Curie European/International Reintegration Grant (303814), and Worldwide Cancer Research project grant (13-0146). J.W. and E.S.P. acknowledge support from Brain Tumor Research and Support across Yorkshire. R.K.M. is a Leeds Cancer Research UK-Centre Clinical Fellow, H.A. B. acknowledges support from EPSRC (EP/M506552/1), C.A. acknowledges support from Cancer Research UK (C48431/A18717; C37059/A1636CRUK).

